# Synergistic innate immune activation and anti-tumor immunity through combined STING and TLR4 stimulation

**DOI:** 10.1101/2024.04.08.588610

**Authors:** Emily F. Higgs, Thomas F. Gajewski

**Affiliations:** Department of Pathology, University of Chicago, Chicago, IL; Department of Medicine, Section of Hematology/Oncology, University of Chicago, Chicago, IL; The Ben May Department for Cancer Research, University of Chicago, Chicago, IL

## Abstract

Previous work has shown that innate immune sensing of tumors involves the host STING pathway, which leads to IFN-β production, dendritic cell (DC) activation, and T cell priming against tumor antigens. This observation has led to the development of STING agonists as a potential cancer therapeutic. However, despite promising results in mouse studies using transplantable tumor models, clinical testing of STING agonists has shown activity in only a minority of patients. Thus, further study of innate immune pathways in anti-tumor immunity is paramount. Innate immune activation in response to a pathogen rarely occurs through stimulation of only one signaling pathway, and activating multiple innate immune pathways similar to a natural infection is one possible strategy to improve the efficacy of STING agonists. To test this, we performed experiments with the STING agonist DMXAA alone or in combination with several TLR agonists. We found that LPS + DMXAA induced significantly greater IFN-β transcription than the sum of either agonist alone. To explain this synergy, we assayed each step of STING pathway signaling. LPS did not increase STING protein aggregation, IRF3 phosphorylation, or IRF3 nuclear translocation beyond what occurred with DMXAA alone. However, since the IFN-β promoter also includes NF-κB binding sites, we additionally examined the NF-κB pathway. In fact, LPS increased the phosphorylation and nuclear translocation of the NF-κB subunit p65, and NF-κB signaling was required for the observed synergy. Intratumoral injection of suboptimal doses of LPS + DMXAA resulted in significantly improved tumor control of B16 melanoma in vivo compared to either agonist alone. Our results suggest that combinatorial signaling through TLR4 and STING results in optimal innate signaling via co-involvement of NF-κB and IRF3, and that combined engagement of these two pathways has therapeutic potential.

## Introduction

Pharmacological engagement of the STING pathway has shown great efficacy in preclinical mouse models, but therapeutic effects in human cancer patients have been limited. The results from early clinical data showing STING agonists alone result in tumor shrinkage in only a subset of patients indicate that deeper knowledge of innate immune pathway activation in response to cancer is needed. In the case of pathogen exposure, innate immune sensing rarely occurs by activating only one signaling pathway, and mammalian organisms have evolved multiple discreet innate immune sensing receptor systems leading to the downstream activation of several transcription factors. The detection of pathogen-associated molecular patterns (PAMPs) or danger-associated molecular patterns (DAMPs) from damaged tissue by specific receptors allows the innate immune system to distinguish between threats and tailor downstream responses accordingly.

A major class of innate receptors for detecting pathogens are the Toll-like receptors (TLRs). TLRs are pattern recognition receptors that reside on the cell surface or in endosomes of various immune cells. They recognize microbial protein or lipid components as well as single or double stranded RNA or unmethylated CpG DNA^1^. TLR signaling leads variably to activation of transcription factors such as NFκB, IRF3, and IRF7, and contributes to the production of proinflammatory cytokines including type I IFNs, TNFα, IL-6, and IL-1β^2^. In addition to cytokine production, TLR activation promotes costimulatory molecule expression on macrophages and DCs, providing an important link between innate and adaptive immune responses.

In the tumor context, IFN-β has been identified as necessary for tumor-specific T cell priming. The IFN-β enhancer contains binding sites for multiple transcription factors, including IRF3, IRF7, NFκB, and AP1, and it is likely that a combination of multiple signaling pathways and downstream transcription factors is necessary for optimal IFN-β transcription^3,4^. In fact, it has been reported that maximal IFN-β transcription is only achieved when all four transcription factor binding sites are present and cells are stimulated with a complex challenge, such as a virus. Stimulation with agonists known to activate these transcription factors individually were unable to elicit strong IFN-β production.

Because the activation of multiple transcription factors can contribute to IFN-β transcription, the IFN-β enhancer is a potential point of interaction between multiple innate immune signaling pathways to impact on IFN-β transcription. This notion led to the idea that a deeper understanding of STING pathway interaction with TLR pathways could inform novel approaches to augment IFN-β production in in the context of cancer. STING agonists alone may result in suboptimal innate immune activation, which could be one reason why many patients do not respond to STING agonist therapy.

Currently there are several TLR agonists in clinical development, and the TLR7 agonist Imiquimod has been FDA approved in basal cell carcinoma among other indications^5^. Additional preclinical and clinical data suggest that achieving successful anti-tumor responses in a large fraction of patients requires combinatorial approaches^6^. We hypothesized that activating multiple signaling pathways may improve innate immune activation in response to tumors and may increase the therapeutic effects of STING agonist therapy. Pursuing rational combinations of STING agonists and TLR agonists has the potential to limit deleterious effects of high dose STING agonist administration and potentially increase the fraction of patients who respond to STING agonist therapy.

## Materials and Methods

### Cell lines and culture conditions

Macrophage and tumor cell lines were passaged in DMEM (Fisher #11995073) supplemented with 10% heat-inactivated fetal bovine serum, 100 U/mL Penicillin/Streptomycin (Fisher 15140122), and 1% Non-Essential Amino Acids (Fisher #11140050). Wild type (WT) mouse macrophages were immortalized as described in Roberson and Walker^7^ and were obtained from the laboratory of Dr. K. Fitzgerald at the University of Massachusetts. Asc^-/-^ macrophages that overexpress STING-HA tag were used to measure STING aggregation. To measure NF-κB activity in reporter macrophages, RAW-Blue™ Cells were used (Invivogen raw-sp). The B16.F10.SIY (henceforth referred to as B16.SIY) melanoma cell line was derived from C57BL/6 mice as described^8^. The DC2.4 dendritic cell line was purchased from Sigma (#SCC142) and cultured in RPMI (Fisher #11875119) supplemented with 10% heat-inactivated fetal bovine serum, 1% L-Glutamine (Fisher #25030081), 1% Non-Essential Amino Acids (Fisher #1140050), 1% HEPES Buffer Solution (Fisher #15630080), and 0.0054% β -Mercaptoethanol (Fisher #21985023).

### In vitro stimulation assays

Cells were seeded in tissue culture-treated 6 well plates at a density of 1 million cells per well. The next day cells were stimulated with one of several agonists depending on the experiment. DMXAA (Cayman Chemical #117570-53-3) and LPS (Cell Signaling #14011) stimulations were performed at 50 μg/mL and 50 ng/mL, respectively, unless otherwise noted. Cells were stimulated for 2 hours at 37° C prior to fixation or lysis, depending on the experiment.

### Quantitative real-time PCR

RNA was extracted from stimulated cells using the Qiagen RNeasy Micro Kit according to the manufacturer’s protocol. Columns were substituted with EconoSpin All-in-one silica membrane mini spin columns (Epoch Life Science). Following isolation, RNA concentration was quantified by nanodrop and diluted to 1.5 μg per reaction with water. RNA was treated with DNAse I (Sigma #4716728001) prior to cDNA synthesis using high-capacity reverse transcriptase (Fisher #4368814). The resulting cDNA was resuspended with water to a final volume of 100 μL, 5 μL of which was used for each qRT-PCR reaction with 20 μL of Taqman gene expression master mix (Fisher #4369514). Roche probes and primers were added to the master mix as described below, and samples were run on a StepOne Plus real-time PCR machine (Applied Biosystems #4376600).

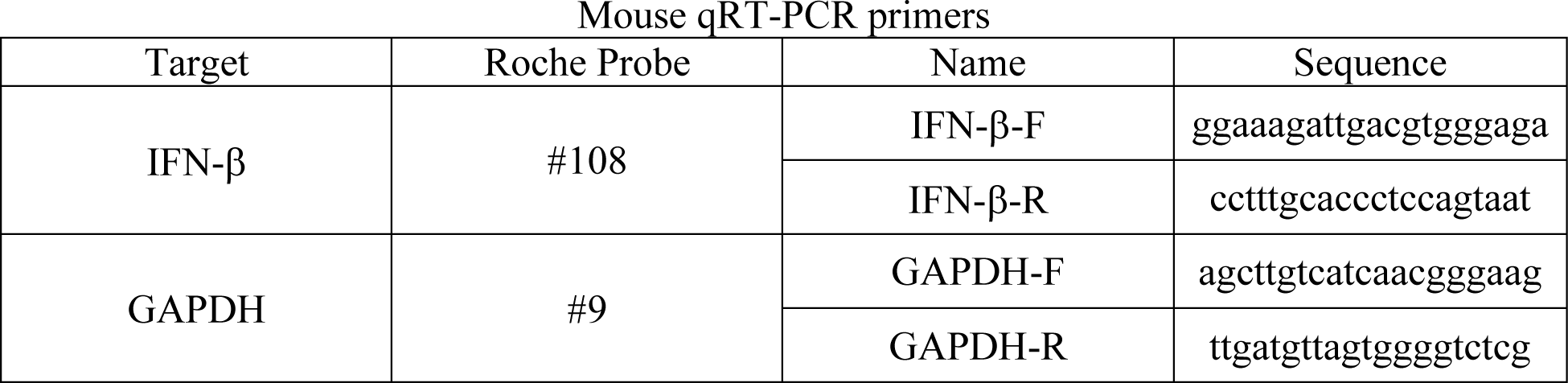

### Immunofluorescent imaging flow cytometry

Macrophages were stimulated at a minimum of 2 million cells per condition. STING aggregation was measured in STING-HA tagged macrophages, whereas IRF3 and p65 nuclear translocation was measured in WT macrophages. Following stimulation, cells were washed with PBS and resuspended in fixation/permeabilization buffer from a kit (Fisher #00552300). The below antibodies were used in intracellular staining for the antigens of interest.

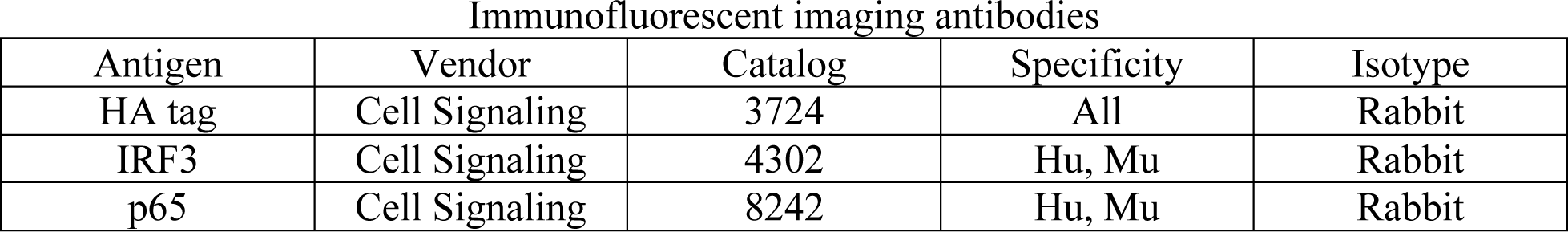

APC-conjugated goat anti-rabbit secondary antibodies were used for secondary staining. Prior to imaging, samples were stained for 5 minutes with 2 drops per mL DAPI (Akoya #FP1490) in permeabilization buffer to label nuclei and washed twice with PBS. An Amnis ImageStreamX machine was used to acquire images, which were then analyzed using IDEAS software according to the manufacturer’s instructions.

### NF-κB reporter assay

NF-κB reporter RAW-Blue™ macrophages were seeded at 50,000 cells per well in a 96 well tissue culture-treated plate. Cells were stimulated the next day with DMXAA, LPS, or both at a range of doses for two hours. Cell suspensions were incubated with QUANTI-Blue™ according to manufacturer’s instructions and SEAP levels were measured using a spectrophotometer at 650 nm.

### Ex vivo stimulation

Spleens from RelA^fl/fl^ x CD11c-Cre mice (Jax 024342 crossed to Jax 008068) were harvested and incubated with digestion buffer for 30 minutes at 37° C while shaking at 200 RPM. The digestion buffer contained RPMI (Fisher #11875119), 2% fetal bovine serum, 200 units/mL bovine pancreas Deoxyribonuclease I (Sigma C5138), 1 mg/mL Hyaluronidase (Sigma H6254), and 1 mg/mL Collagenase Type IV (Sigma C5138). Digested spleens were then mashed through a 70 μm cell strainer and washed with 15 mL PBS. Gey’s solution (500 mL water with 4.15 g NH_4_Cl plus 0.5 g KHCO_3_ and filter sterilized) was added for 1 minute to lyse red blood cells in the mixture. CD11c^+^ cells were isolated using the CD11c MicroBeads UltraPure kit using LS columns from Miltenyi (#130125835) according to manufacturer’s instructions. Purified cells from each spleen were resuspended in complete DMEM, divided into four groups, and stimulated for two hours with the conditions control, LPS, DMXAA, or LPS + DMXAA. Following stimulation, qRT-PCR for IFN-β and GAPDH was performed. Cre-negative spleens from littermate control mice were used as controls. Approximately 1/50^th^ of each sample was removed following CD11c isolation to check the cellular purity by flow cytometry, and only samples containing 5% or fewer of T, B, and NK cells were used. The flow cytometry panel used is described below.

| Marker | Clone | Fluorophore | Vendor |
| --- | --- | --- | --- |
| B220 | RA3-6B2 | FITC | PharMingen |
| CD11b | M1/70 | BV711 | BioLegend |
| CD11c | N418 | PE | BioLegend |
| CD3 | 17A2 | AF700 | BioLegend |
| CD45 | 30-F11 | APC-Cy7 | BioLegend |
| I-A/I-E | M5/114.15.2 | PacBlue | BioLegend |
| GR-1 | RB6-8C5 | PE-Cy7 | BioLegend |
| Viability | Fixable Viability Dye eFluor™ 506 |  | eBioscience |

### Transplantable tumor model

C57BL/6 obtained from Taconic Biosciences (Hudson, NY) were housed in specific pathogen-free (SPF) conditions. TLR4 knockout (Jax 029015) and STING knockout (Jax 025805) mice were also housed in SPF conditions. Mice were cohoused and given 1 million live B16.SIY tumor cells at 6-8 weeks of age via subcutaneous injection. Tumor volume was calculated by tumor length x width x height as measured with calipers three times per week until endpoint size was reached. In vivo DMXAA injections were performed intratumorally with a single dose of 250 μg in 50 μL per mouse unless otherwise specified. In vivo LPS injections were performed intratumorally with a single dose of 250 ng in 50 μL per mouse unless otherwise specified. Both LPS and DMXAA were injected on day 11 post tumor inoculation.

### Statistical analysis

Tumor growth curves were analyzed in GraphPad PRISM. Differences in growth were determined using Tukey’s multiple comparisons post-test. For other comparisons between two groups, unpaired, two-tailed Student’s tests were used. For comparisons between three or more groups, one-way ANOVA’s were used to evaluate differences, with Benjamini-Hochberg FDR correction for multiple comparisons^9^. Statistical significance was considered to be p < 0.05 and was denoted as follows: *p < 0.05, **p < 0.01, ***p < 0.001, ****p < 0.0001. All statistical tests were performed using GraphPad PRISM and R.

## Results

### LPS synergizes with DMXAA in vitro

We initially examined several TLR agonists, including Pam3CSK4 (TLR1/2), Poly:IC (TLR3), LPS (TLR4), Gardiquimod (TLR7), and CpG ODN1668 (TLR9) for their ability to induce IFN-β transcription in macrophages. We found that each TLR agonist tested individually at a range of concentrations induced far less IFN-β production than the STING agonist DMXAA (Figure 1A). However, when tested in combination with DMXAA, LPS induced a synergistic increase in IFN-β transcription, which was significantly greater than the sum of induction by either agonist alone (Figure 1B). This was true across a range of doses tested, and LPS significantly increased the amount of IFN-β produced in response to low doses of DMXAA (Figure 1C). The other TLR agonists in combination with DMXAA did not significantly increase IFN-β transcription compared to either agonist alone. The synergistic effect of LPS and DMXAA was also observed in the DC2.4 cell line, indicating that the synergy observed is not a macrophage-restricted phenomenon but also occurs in DCs (Figure 1D).

**FIGURE 1.**
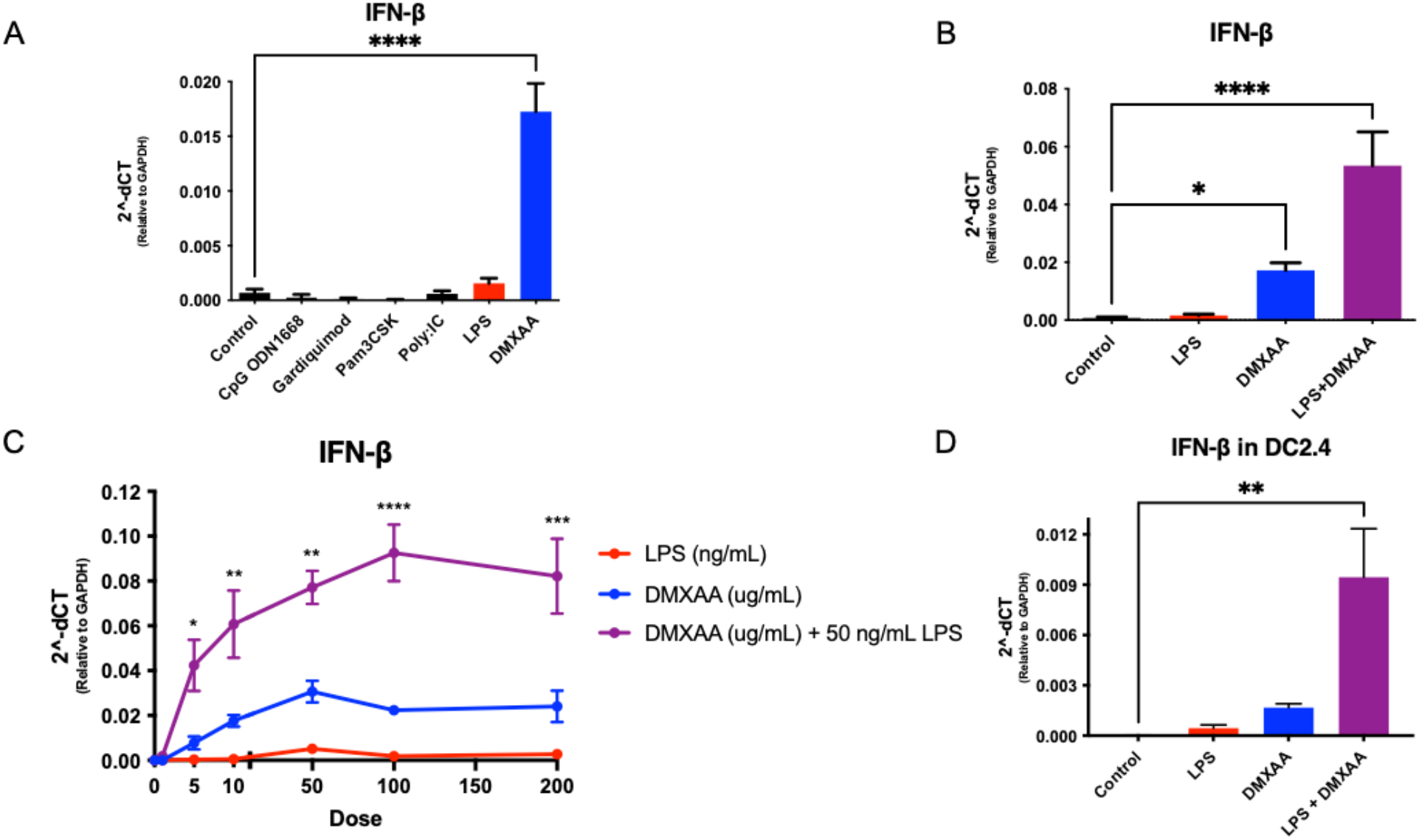
LPS and STING agonist DMXAA synergize to induce IFN-β transcription. (A) IFN-β transcription following stimulation of macrophages in vitro with TLR agonists CpG ODN 1668, Gardiquimod, Pam3CSK, Poly:IC, LPS, and STING agonist DMXAA. (**B**) IFN-β transcription following stimulation of macrophages in vitro with LPS, DMXAA, or combination. (**C**) IFN-β transcription following stimulation of macrophages in vitro with dose titrations of LPS, DMXAA, or DMXAA + 50 ng/mL LPS. (**D**) IFN-β transcription following stimulation of DC2.4 cells in vitro with LPS, DMXAA, or combination.

Thus, LPS was able to significantly increase the maximal amount of IFN-β produced in response to DMXAA. Even low doses of DMXAA combined with LPS induced greater IFN-β than high doses of DMXAA alone.

### LPS does not augment DMXAA signaling through STING or IRF3 activation

Next, we sought to determine the mechanism by which the synergy between LPS and DMXAA was occurring, and first focused on the core STING pathway itself. Downstream signaling following STING engagement is well-characterized and involves STING aggregation followed by TBK1 and IRF3 phosphorylation. This then leads to IRF3 nuclear translocation where it can carry out its transcriptional activity and induce IFN-β gene expression. To assess STING aggregation in response to DMXAA, LPS, and the combination, we stimulated STING-HA labeled macrophages and performed ImageStream analysis to visualize STING aggregation. Robust STING activation was induced by DMXAA and was also present in the LPS+DMXAA condition, but not in the condition with LPS alone (Figure 2A, Figure 2B). Similarly, LPS did not significantly induce IRF3 nuclear translocation or increase the nuclear translocation of IRF3 induced by DMXAA alone (Figure 2C, Figure 2D).

**FIGURE 2.**
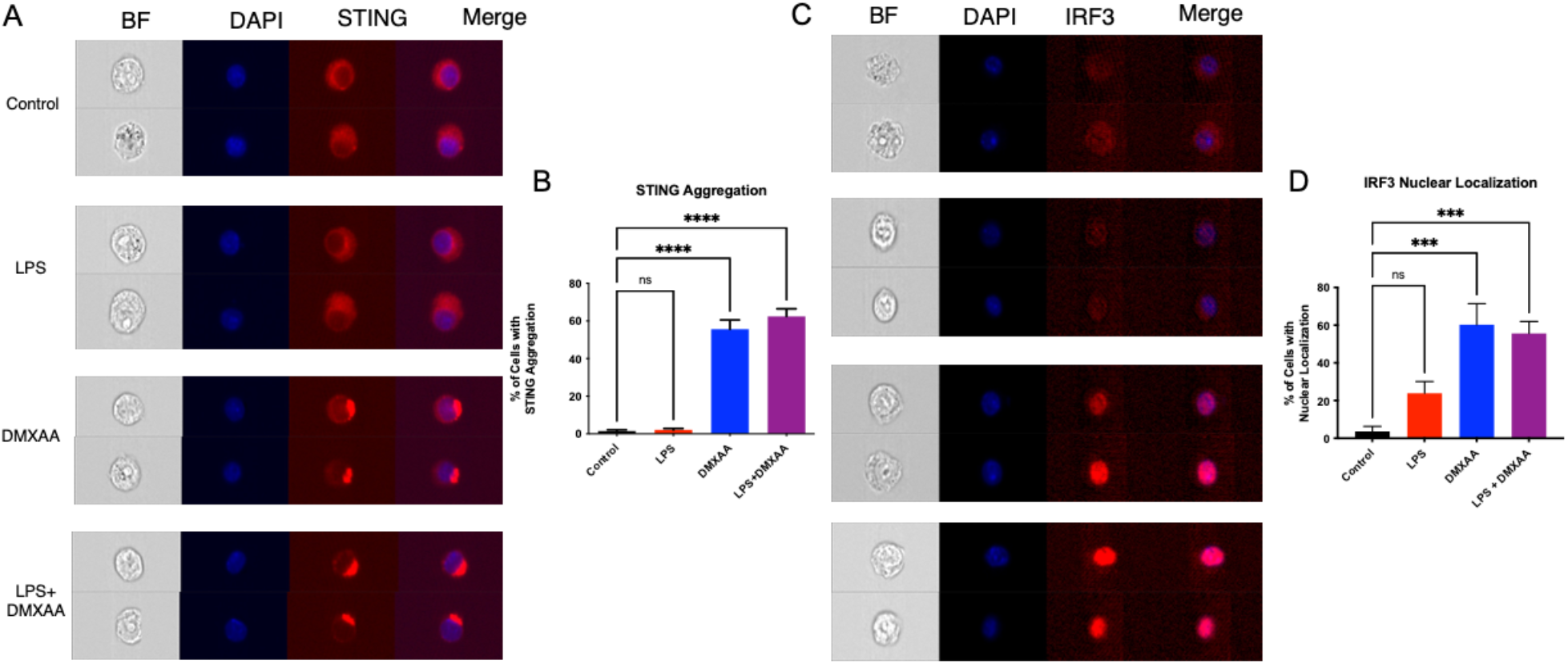
LPS does not strongly affect STING aggregation or IRF3 nuclear translocation. (**A**) ImageStream images of STING aggregation in STING-HA macrophages stimulated with LPS, DMXAA, LPS + DMXAA. (**B**) Quantification of percentage of cells with STING aggregation following stimulation. (**C**) ImageStream images of IRF3 nuclear localization in macrophages stimulated with LPS, DMXAA, LPS + DMXAA. (**D**) Quantification of percentage of cells with nuclear localization based on overlap with DAPI.

These data suggest that LPS is not augmenting the IFN-β response to DMXAA by affecting STING directly or the transcription factor IRF3.

### LPS signaling through NFκB is required for synergy

Since signaling downstream of LPS is known to activate NFκB, and NFκB can contribute to transcriptional regulation of the IFN-β promoter/enhancer, we next examined NFκB in response to stimulation with these agonists. To study NFκB activity, we used ImageStream to measure nuclear translocation of the NFκB subunit p65. Although LPS failed to induce IRF3 nuclear translocation, LPS induced robust p65 nuclear translocation (Figure 3A, Figure 3B). DMXAA had a relatively minor effect on p65 nuclear translocation, and the combination of LPS + DMXAA appeared similar to LPS alone. Next, we examined NFκB activity directly using RAW-Blue™ reporter macrophages. Across a range of doses, LPS induced significantly greater reporter activity than did DMXAA, and the LPS + DMXAA condition appeared similar to LPS alone (Figure 3C).

**FIGURE 3.**
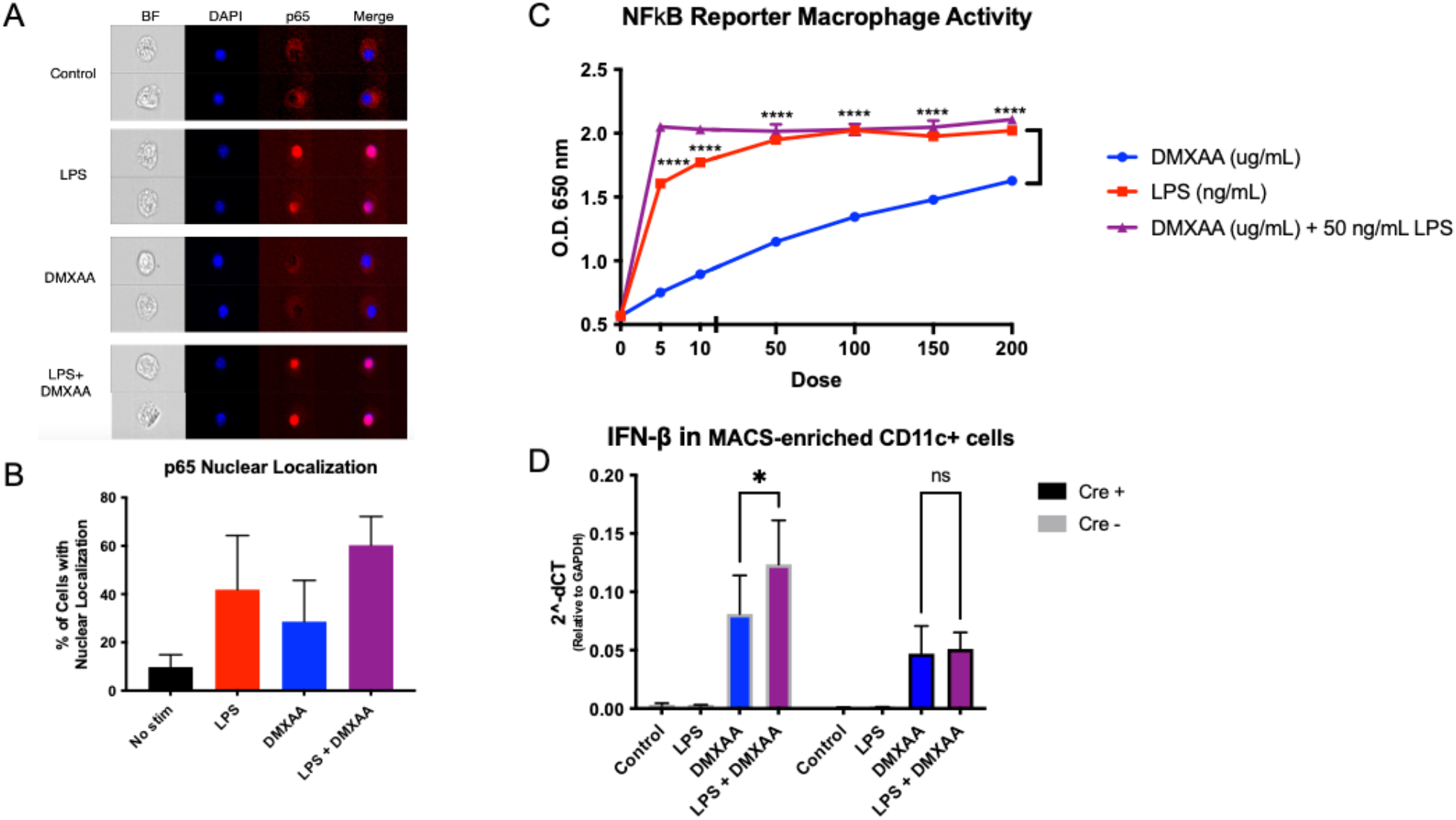
LPS promotes NFκB activation by increasing p65 nuclear translocation. (**A**) ImageStream images of p65 nuclear localization in macrophages stimulated with LPS, DMXAA, LPS + DMXAA. (**B**) Quantification of percentage of cells with nuclear localization based on overlap with DAPI. (**C**) NFκB reporter activity in RAW-Blue™ macrophages stimulated with following stimulation with dose titrations of LPS, DMXAA, or DMXAA + 50 ng/mL LPS. (**D**) IFN-β transcription in CD11c^+^ cells purified from RelA x CD11c-Cre^+^ and RelA x CD11c-Cre^-^ mice stimulated with LPS, DMXAA, LPS + DMXAA.

Since NFκB was readily induced by LPS stimulation, we examined whether it was required for the synergistic induction of IFN-β expression. To test the necessity of NFκB for the synergy between LPS and DMXAA, we isolated CD11c^+^ cells from p65^fl/fl^ mice that were CD11c-Cre^+^ or CD11c-Cre^-^ and measured IFN-β following agonist stimulation. As we had observed previously with macrophages and the DC cell line, CD11c^+^ cells from Cre^-^ mice that had p65 intact showed increased IFN-β transcription with LPS+DMXAA compared to DMXAA stimulation alone (Figure 3D). However, with CD11c^+^ cells deleted of p65, no such increased was observed with combination treatment. This result indicates that p65 is necessary for the synergy between LPS and DMXAA in causing increased IFN-β production.

Overall, these data support our model that DMXAA primarily signals through the transcription factor IRF3, and LPS primarily signals through NFκB. Activation of both of these transcription factors with agonists for both pathways induces synergistic levels of IFN-β transcription.

### LPS synergizes with DMXAA to promote tumor control in vivo

Because LPS + DMXAA induced such a strong IFN-β response in vitro, we next wanted to test if LPS could promote tumor control in vivo with DMXAA. To do so, we used a suboptimal 250 ug dose of DMXAA, and performed a single intratumoral injection with or without 250 ng LPS. As evidenced by the tumor growth curves, both LPS and DMXAA alone resulted in modest tumor control. However, there was a significant improvement when both agonists were administered together (Figure 4).

**FIGURE 4.**
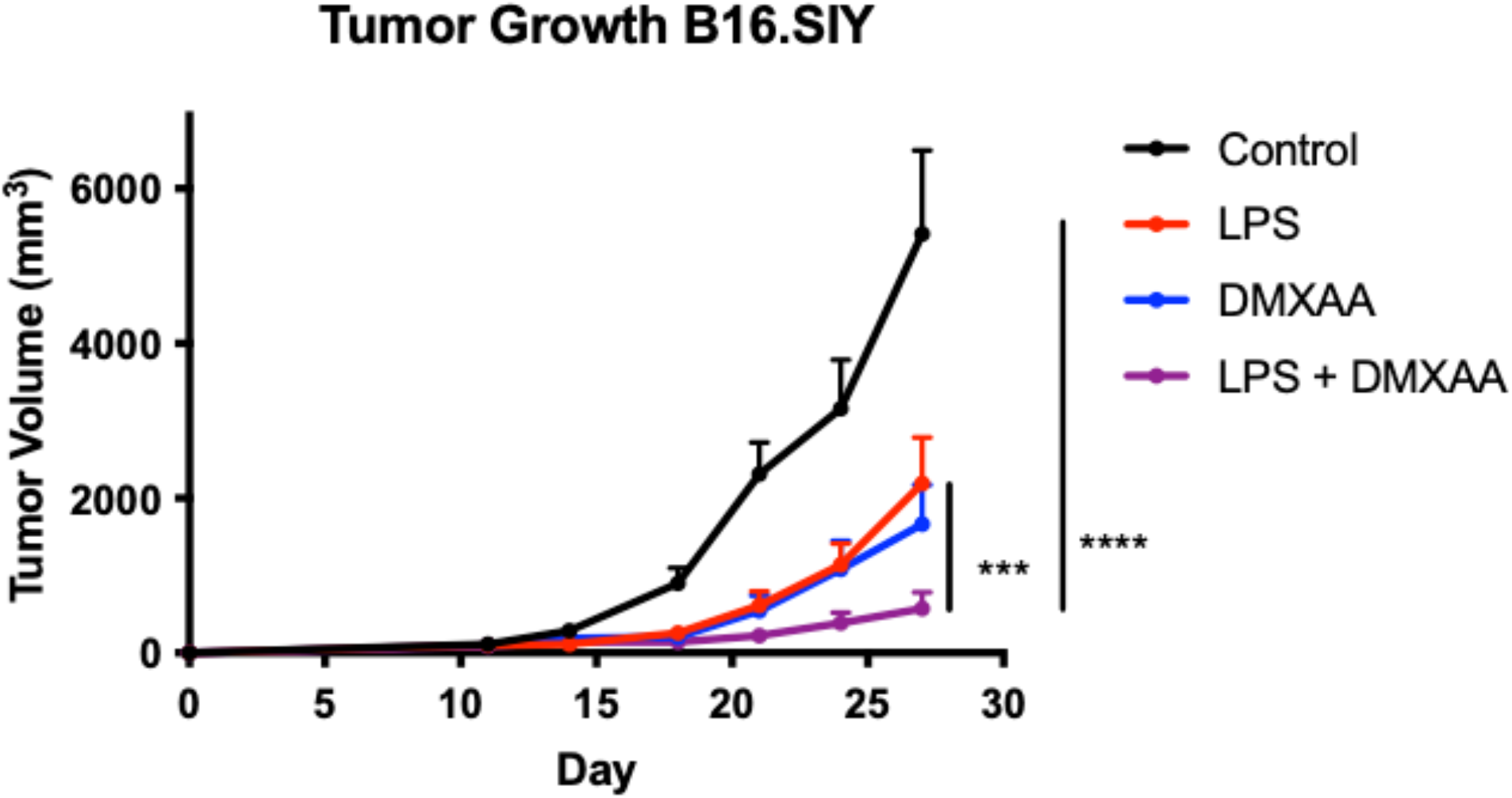
Intratumoral LPS improves tumor control alone and in combination with DMXAA. B16.SIY tumor growth in C57BL/6 mice treated intratumorally on day 11 with 250 μg DMXAA, 250 ng LPS, or both.

This observation indicates that combining agonists for different innate immune pathways, specifically the STING and TLR4 pathways, can result in improved anti-tumor immune responses. To determine whether host expression of STING and TLR4 were required for this improved tumor control, the same in vivo injection strategy was performed in STING^-/-^ and TLR4^-/-^ mice. LPS showed some anti-tumor activity in STING^-/-^ mice, but there was no additional benefit with DMXAA in these mice (Figure 5A). Similarly, DMXAA showed some activity in TLR4^-/-^ mice, but synergy was not observed between the two agonists in these mice either (Figure 5B).

**FIGURE 5.**
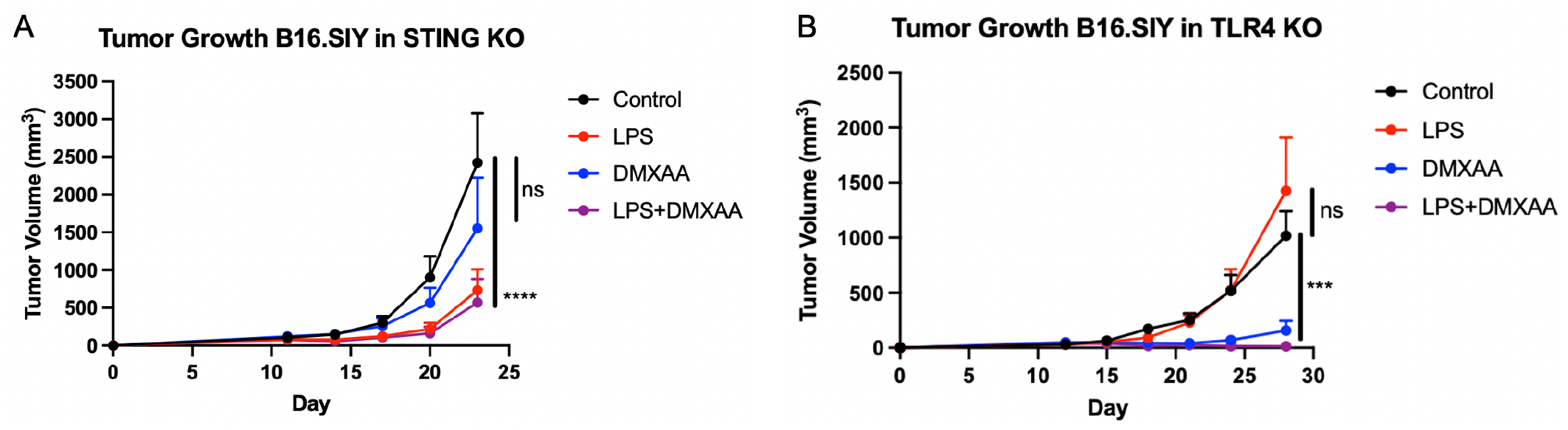
In vivo benefit of DMXAA and LPS combination is dependent on STING and TLR4. (**A**) B16.SIY tumor growth in STING^-/-^ mice treated intratumorally on day 11 with 250 μg DMXAA, 250 ng LPS, or both. (**B**) B16.SIY tumor growth in TLR4^-/-^ mice treated intratumorally on day 11 with 250 μg DMXAA, 250 ng LPS, or both.

These data support the notion that host STING signaling and TLR4 signaling must both be intact in order to observe improved tumor control in response to LPS + DMXAA therapy.

## Discussion

In this study, we examined the hypothesis that in some cases activating the STING pathway alone is insufficient to optimally induce IFN-β transcription, and that using a combination of innate immune agonists is superior. By treating APCs with a range of concentrations of STING agonist in addition to TLR agonists, we determined LPS was capable of synergizing with the STING pathway for optimal IFN-β production that is greater than either agonist is capable of inducing alone. We then interrogated downstream signaling events to determine the mechanism driving additional IFN-β production. Based on in vitro experiments performed in macrophages and dendritic cells, DMXAA was found to signal primarily through the transcription factor IRF3, whereas LPS signaled primarily through the transcription factor NFκB. Together, these data suggest a working model that agonists targeting both of these pathways induce better activation of both IRF3 and NFκB, resulting in augmented IFN-β transcription by innate immune cells and superior tumor control (Figure 6).

**FIGURE 6.**
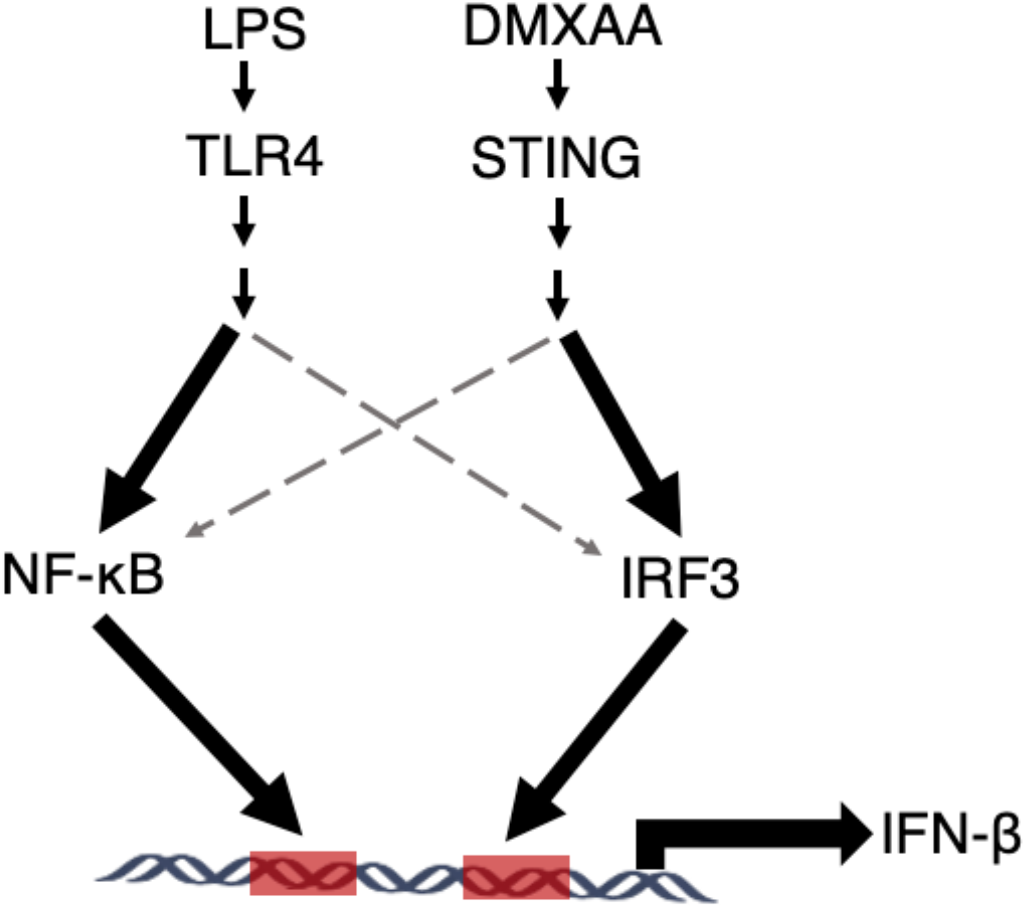
Working model of synergy between LPS and DMXAA.

Our current working model is that LPS primarily signals through TLR4 to activate the transcription factor NFκB, while DMXAA signaling through STING primarily activates the transcription factor IRF3. When both agonists are present to activate both transcription factors, synergistic levels of IFN-β are produced.

IRF3 activation downstream of STING has been well-characterized, and most studies of STING signaling have focused on this arm of the pathway^10–12^. More recently however, there has been some evidence implicating a role for NFκB as well^13^. Three distant “Alu” sites associate with the IFN-β locus and are bound by NFκB prior to it binding the IFN-β promotor^14^. While further characterization is needed, it is thought that NFκB binding to these “Alu” sites may epigenetically open the IFN-β promotor region and allow the other transcription factors to bind. Our results indicate that NFκB activation downstream of STING agonist treatment is significantly less than NFκB activation downstream of LPS treatment. This could help explain why LPS is able to dramatically augment IFN-β transcription, if NFκB is required for optimal IFN-β locus accessibility.

Another important consideration is that not every cell produces IFN-β following stimulation, and that proper transcription factor binding leading to transcription is a stochastic process. Evidence for this comes from the fact that single cell cloning from a pool of cells capable of expressing IFN-β produces pools of cells that express IFN-β at the same proportion^15^. This could be due to the fact that there is a limited supply of the transcription factors binding to IFN-β, and that there may not be sufficient quantities of each free to bind the IFN-β in every cell. Consistent with this hypothesis, overexpression of either IRF3 or NFκB results in a higher proportion of cells that express IFN-β^16,17^. This raises an interesting future question as to whether LPS stimulation drives more IFN-β production per cell or whether it increases the number of cells that produce IFN-β.

Additionally, LPS promoted tumor control in vivo when combined with low dose STING agonist administration. The ability to achieve a robust anti-tumor immune response with lower STING agonist doses may protect against some of the negative effects of STING agonist therapy. Treatment with high-dose STING agonist has been shown to actually result in reduced tumor-specific T cell priming and can even be cytotoxic to T cells, as T cell-intrinsic STING signaling can promote apoptosis. Therefore, this combination may represent a viable therapeutic strategy to improve STING agonist response rates and mitigate some of the negative effects associated with higher doses of STING agonist therapy.

## Acknowledgments

We would like to thank the University of Chicago Flow Cytometry Core Facility and Animal Resources Center.

